# Quantifying the impact of sample, instrument, and data processing on biological signatures detected with Raman spectroscopy

**DOI:** 10.1101/2023.06.01.543279

**Authors:** Jasmina Wiemann, Philipp R. Heck

## Abstract

Raman spectroscopy is a popular tool for characterizing complex biological materials and their geological remains^1-10^. Ordination methods, such as Principal Component Analysis (PCA), rely on spectral variance to create a compositional space^1^, the ChemoSpace, grouping samples based on spectroscopic manifestations that reflect different biological properties or geological processes^1-7^. PCA allows to reduce the dimensionality of complex spectroscopic data and facilitates the extraction of relevant informative features into data formats suitable for downstream statistical analyses, thus representing an essential first step in the development of diagnostic biosignatures. However, there is presently no systematic survey of the impact of sample, instrument, and spectral processing on the occupation of the ChemoSpace. Here the influence of sample count, signal-to-noise ratios, spectrometer decalibration, baseline subtraction routines, and spectral normalization on ChemoSpace grouping is investigated using synthetic spectra. Increase in sample size improves the dissociation of sample groups in the ChemoSpace, however, a stable pattern in occupation can be achieved with less than 10 samples per group. Systemic noise of different amplitude and frequency, features that can be introduced by instrument or sample^11,12^, are eliminated by PCA even when spectra of differing signal-to-noise ratios are compared. Routine offsets (± 1 cm^−1^) in spectrometer calibration contribute to less than 0.1% of the total spectral variance captured in the ChemoSpace, and do not obscure biological information. Standard adaptive baselining, together with normalization, increase spectral comparability and facilitate the extraction of informative features. The ChemoSpace approach to biosignatures represents a powerful tool for exploring, denoising, and integrating molecular biological information from modern and ancient organismal samples.

## 1. INTRODUCTION

Raman spectroscopy allows non-destructive compositional fingerprinting of complex biological and geological materials^1-10^. Rapidly generated *in situ* spectra yield information on covalent, ionic, and non-covalent bioinorganic interactions enabling a comparative search for informative heterogeneities across a diversity of samples^1^, such as modern organismal tissues and their fossilization products. Spectroscopic biosignatures, such as phylogenetic and metabolic signals, represent diagnostic tools in cancer research^3-7^, and a number of signatures present in fresh tissues preserve, occasionally altered but not unrecognizable, in fossilized carbonaceous tissues: In integrative data sets, spectroscopic signatures reflecting the relative abundance of different organic functional groups^1^ and organo-mineral interactions^2^ encode molecular manifestations of phylogenetic affinity^2-7,13-15^, physiology^2-7,13-19^, and degree and mode of environmental or diagenetic alteration^1-2,20^. These signals are relative and can only be analyzed in a comparative framework^1-7,13-20^.

Spectra collected across a diversity of samples may contain additional unwanted signals that reduce the signal-to-noise ratio^1,2^. Examples include a non-linear background based on sample fluorescence induced by the excitation source^1,12^, lower intensity counts due to diffusive scattering at rough sample surfaces^1,12^, and (quasi-)sinusoidal noise resulting from reflective scattering at layers with different optical properties within a sample or introduced by certain laser-cancelling filters in combination with specific line gratings^11,12^. Most of these unwanted signals are expressed as spectral baseline functions of different periodicity, amplitude, and frequency^1,11,12^. Noisy spectra are a well-known challenge in biological spectroscopy^1,7^ and processing routines, including adaptive baselining (background correction sensitive to the total spectral curve) and normalization (spectral intensity scaling based on individual peaks or integrated areas), are employed to minimize the impact of unwanted signals on data interpretation^1,2,7,12^. Similarly, spectral phase shift introduced by temperature-based instrument decalibration can be traced and corrected between analytical sessions^21^ but non-linear decalibration rates render correction (up to ± 1 cm^-1^ wavenumbers) during an analytical session challenging.

In the last 30 years, spectroscopy has shifted from exclusively qualitative interpretation of Raman spectral data^8-10^ toward a comparative approach^1,2,5-7^ that relies, as an essential first step in the data analysis, on ordination methods (dimensionality reduction) such as Principal Component Analysis (PCA). PCA allows to explore, denoise (Fig. 1), identify, and extract informative heterogeneities (effectively ‘latent variables’) from sets of inherently complex spectra, each characterized by a very large number of data points collected over the wavenumber range^1,2,7,13-20^. PCA captures the covariance of spectral features in an *n*-dimensional compositional space, the ChemoSpace, where *n* equals the number of features considered^22,23^. The ChemoSpace is based on a variance-covariance matrix ([*number of spectra*] x [*number of features*])^22,23^.

**Figure 1:**
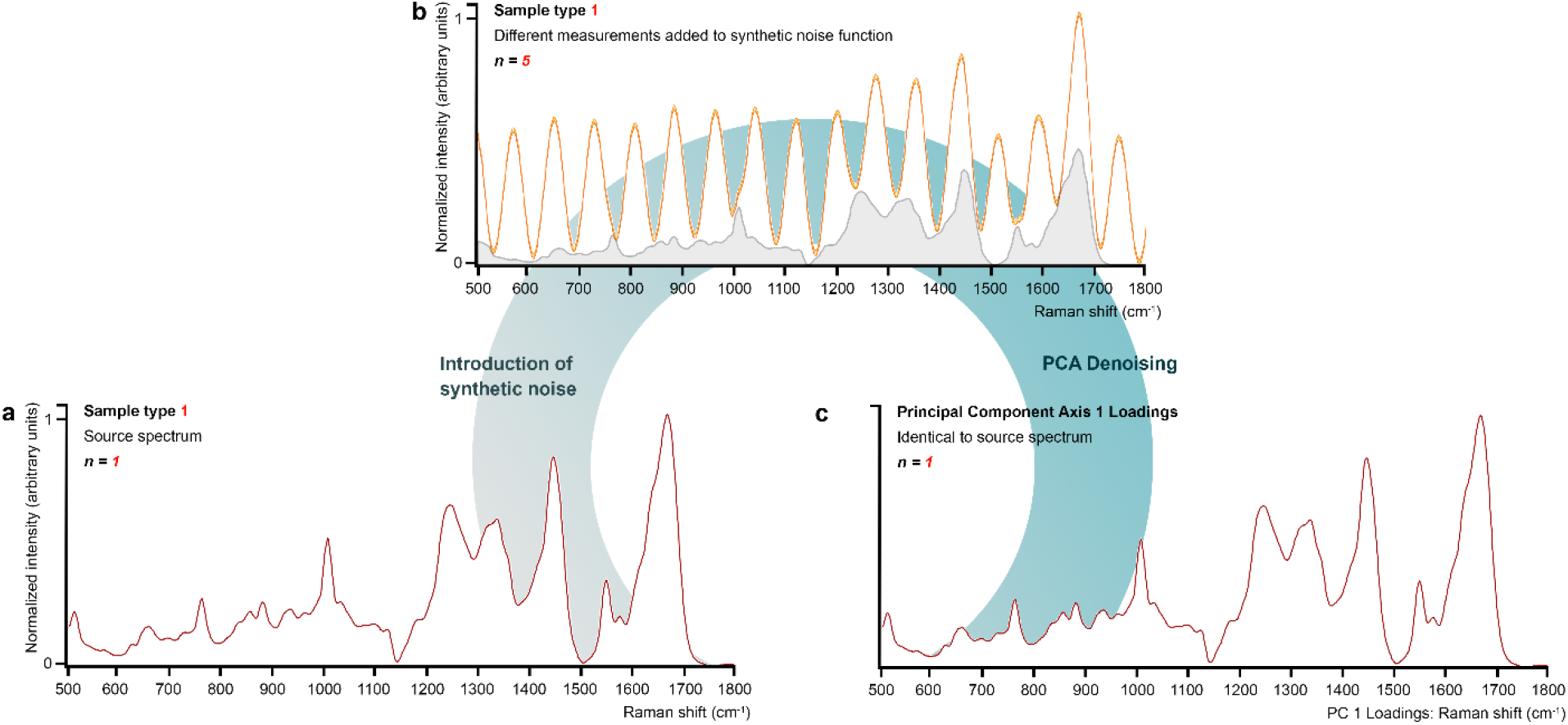
Schematic illustration showcasing the denoising potential of Principal Component Analysis (PCA). **a** n=1 Raman spectrum of the sample type 1 plotted over the organic fingerprint region. A set (n=5) of synthetic technical replicates based on the spectrum plotted in **a** were summed with the high-frequency synthetic noise functions shown in Fig. 3a, in order to generate the n=5 spectra shown in **b. b** The artificially noised n=5 varieties of the source spectrum shown in **a**. The source spectrum in **a** is plotted under the summed spectral curves and is shaded in grey. PCA is applied to the n=5 artifically noised spectra. **c** Resulting PC 1 axis loadings plotted over the organic fingerprint region match the original source spectrum shown in **a**. All artificial noise has been removed by PCA. PCA allows for the robust denoising of spectroscopic data collected for biological or paleontological samples.

The general order of magnitude of the minimum number of spectra required to achieve a stable pattern in the Raman ChemoSpace occupation remains yet to be determined: the data point distribution across the ChemoSpace changes with the number and type of spectra or selected peaks included in the analysis – increasing the number of considered spectra increases the statistical power of sample group separation. Once the number of spectra included is representative of compositional diversity in a sample set, a stable pattern in ChemoSpace occupation is reached.

The number of considered features and thus dimensions of the ChemoSpace (*n*) can encompass all the data points that contribute to a spectrum or, more commonly, selected peaks of interest^1,2^. PCA benefits from the subsampling of prominent peaks in spectra, an approach that prevents overweighting broad signals and uninformative spectral regions^24^. Because the normalized intensities of Raman peaks represent the relative abundancies of molecular features, the ChemoSpace can be thought of as a multivariate application of the Beer-Lambert law, which defines that the spectroscopic signal of a compound is mathematically linked to its abundance in the sample. When spectra of different biological sample types are analyzed by means of PCA, variance corresponding to different biosignatures is commonly expressed by the first two or three principal components (PCs) – the axes of the ChemoSpace displaying different aspects of variance in the data, sorted by descending contributions to the total variance^1,2,23^. Based on the information represented along the PCs and the distribution of eigenvectors which illustrate the impact of individual peaks on the placing of a sample in the ChemoSpace, PCA allows for the exploration and identification of features that are particularly informative for sample grouping, and thus represents an essential tool for the subsampling of spectral data points required toward downstream classification or cluster analyses.

The impact of analytical variables and different types of unwanted spectral features on classification approaches to spectroscopic biosignatures, such as Linear Discriminant Analysis (LDA)^25^ and its corresponding machine-learning tool (Support Vector Machine, SVM)^26,27^ is known and has led to a number of end user recommendations^25-27^, but it is only incompletely characterized for the PCA ChemoSpace. Given the potential of the ChemoSpace to address questions in modern biology^1,3-4^, clinical diagnostics^5-7^, paleontology^2,13-20^, geology^28^, and astrobiology^28^, a systematic survey of the impact of sample size (Fig. 2), spectral signal-to-noise ratios (Figs. 3, 4a, c, 5), spectrometer decalibration (Fig. 6), baseline subtraction routines (Fig. 7), and normalization procedures (Fig. 4b, d) on informative ChemoSpace grouping is overdue.

**Figure 2:**
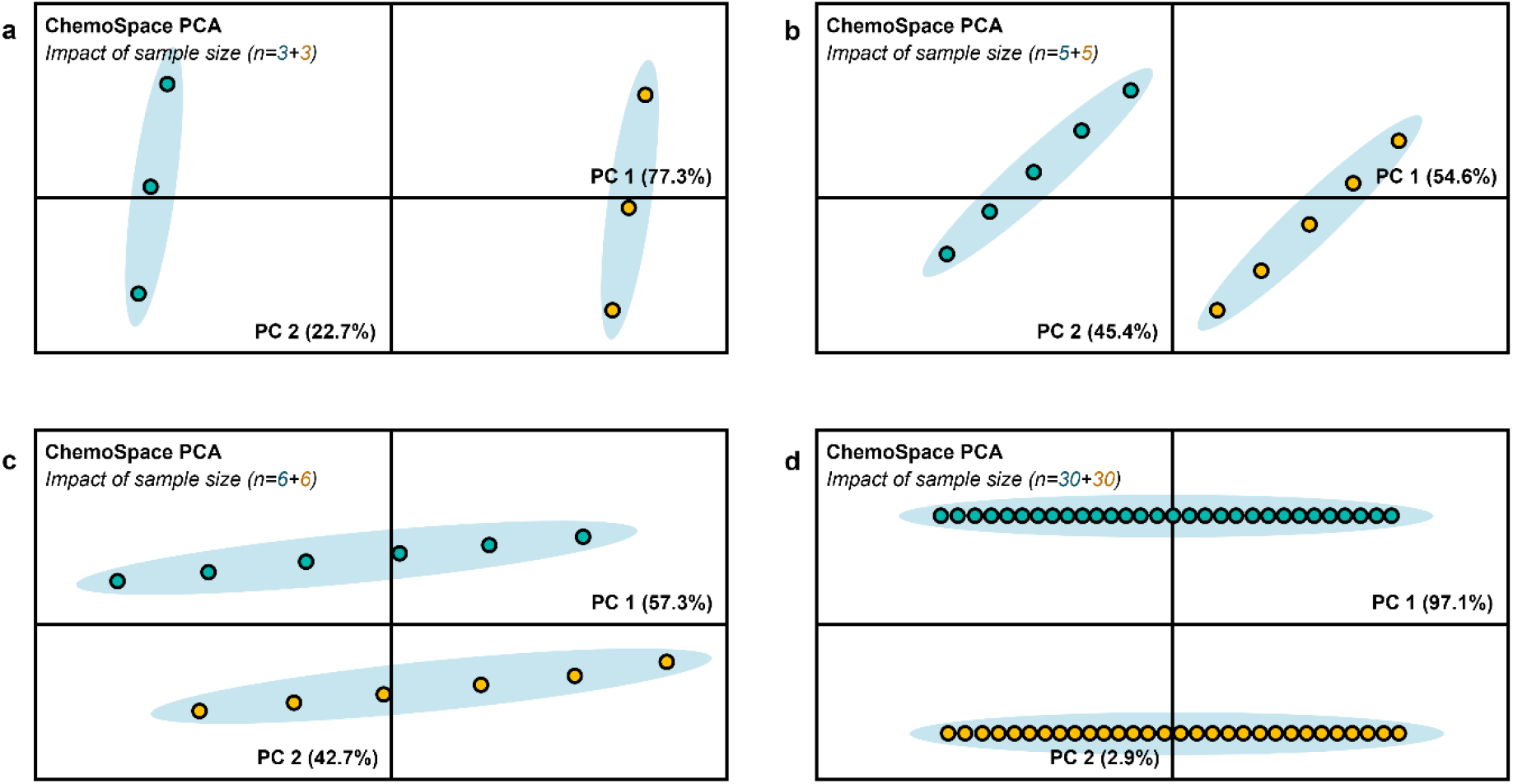
The impact of sample size on the ChemoSpace occupation. **a** ChemoSpace plot resulting from n=3 varieties (synthetic technical replicates) of two sample types (1: teal; 2: orange). **b** ChemoSpace plot resulting from n=5 varieties (synthetic technical replicates) of the two sample types (1: teal; 2: orange). **c** ChemoSpace plot resulting from n=6 varieties (synthetic technical replicates) of the two sample types (1: teal; 2: orange). **d** ChemoSpace plot resulting from n=30 varieties (synthetic technical replicates) of the two sample types (1: teal; 2: orange). All PC loadings are listed in the ChemoSpace plots.

**Figure 3:**
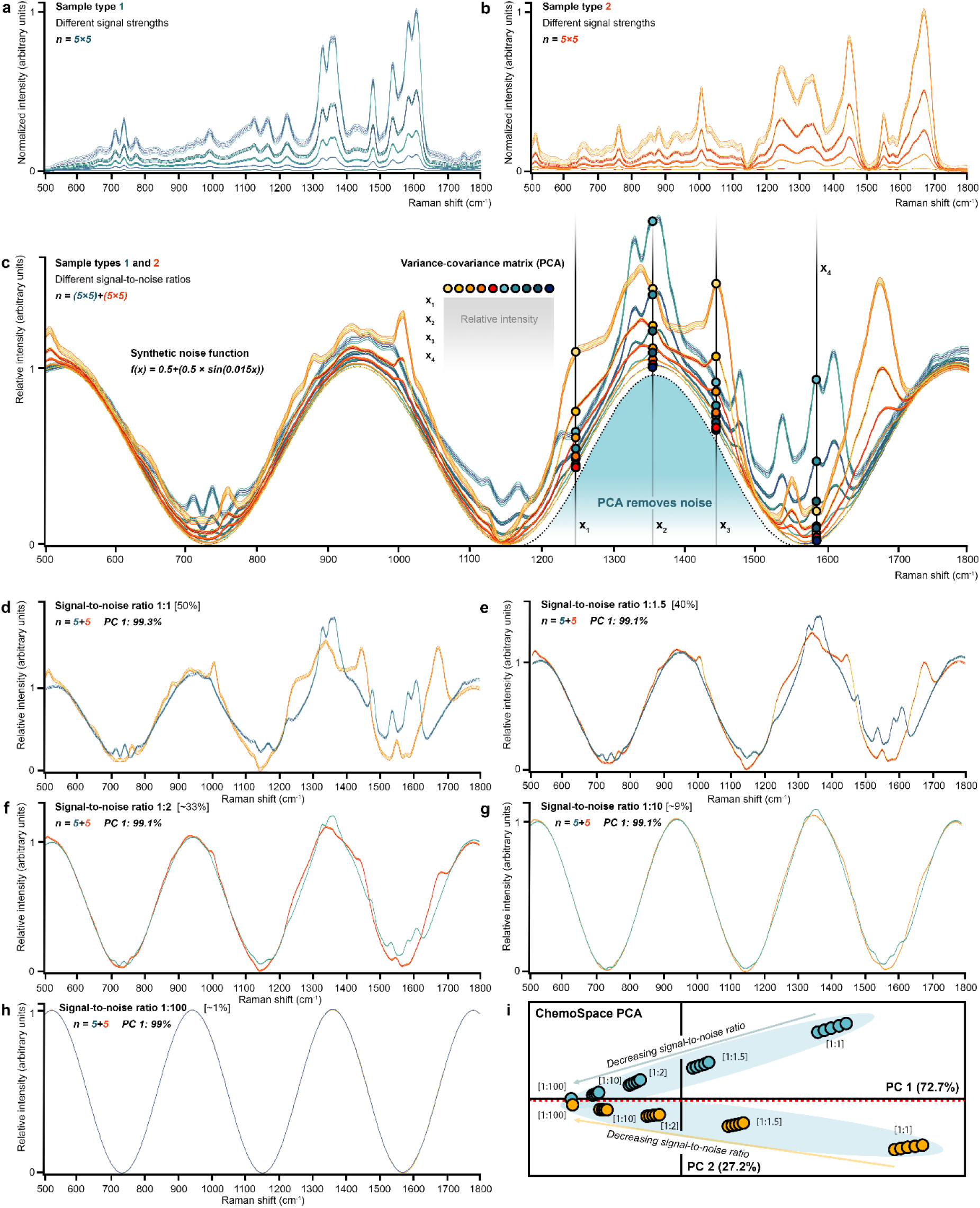
The impact of systemic low-frequency sinusoidal noise and different signal-to-noise ratios on ChemoSpace occupation. **a** Plot of n=5 varieties (synthetic technical replicates) of n=5 differently scaled sets of spectra corresponding to the sample type 1 over the organic fingerprint region. **b** Plot of n=5 varieties (synthetic technical replicates) of n=5 differently scaled sets of spectra corresponding to the sample type 2 over the organic fingerprint region. **c** Plot of n=5 varieties (synthetic technical replicates) of n=5 differently scaled sets of spectra corresponding to the two sample types (1: teal hues; 2: orange hues) added to the normalized synthetic noise function (for details see figure or Methods) over the organic fingerprint region. Four Raman band positions are indicated (x_1_ – x_4_) and the colored data points label the mean average intensity of the individual sets of spectra, in order to visually explain how the variance-covariance matrix is built. Signal-to-noise ratios ranging from ∼1 – 50%. Sets of spectra matching in their signal-to-noise ratio are extracted in **d-h. d** Set of spectra extracted from **c** with a signal-to-noise-ratio of 1:1. **e** Set of spectra extracted from **c** with a signal-to-noise-ratio of 1:1.5. **f** Set of spectra extracted from **c** with a signal-to-noise-ratio of 1:2. **g** Set of spectra extracted from **c** with a signal-to-noise-ratio of 1:10. **h** Set of spectra extracted from **c** with a signal-to-noise-ratio of 1:100. **i** ChemoSpace across PCs 1 and 2 based on a variance-covariance matrix including select relative intensities (see Methods) extracted from the plot in **c**. Data point fill colors correspond to the sample type (1: teal hues; 2: orange hues; compare **c**). The different signal-to-noise ratios are shown for groups of data points, and the labelled arrows indicate general patterns in the data dristribution across the ChemoSpace.

**Figure 4:**
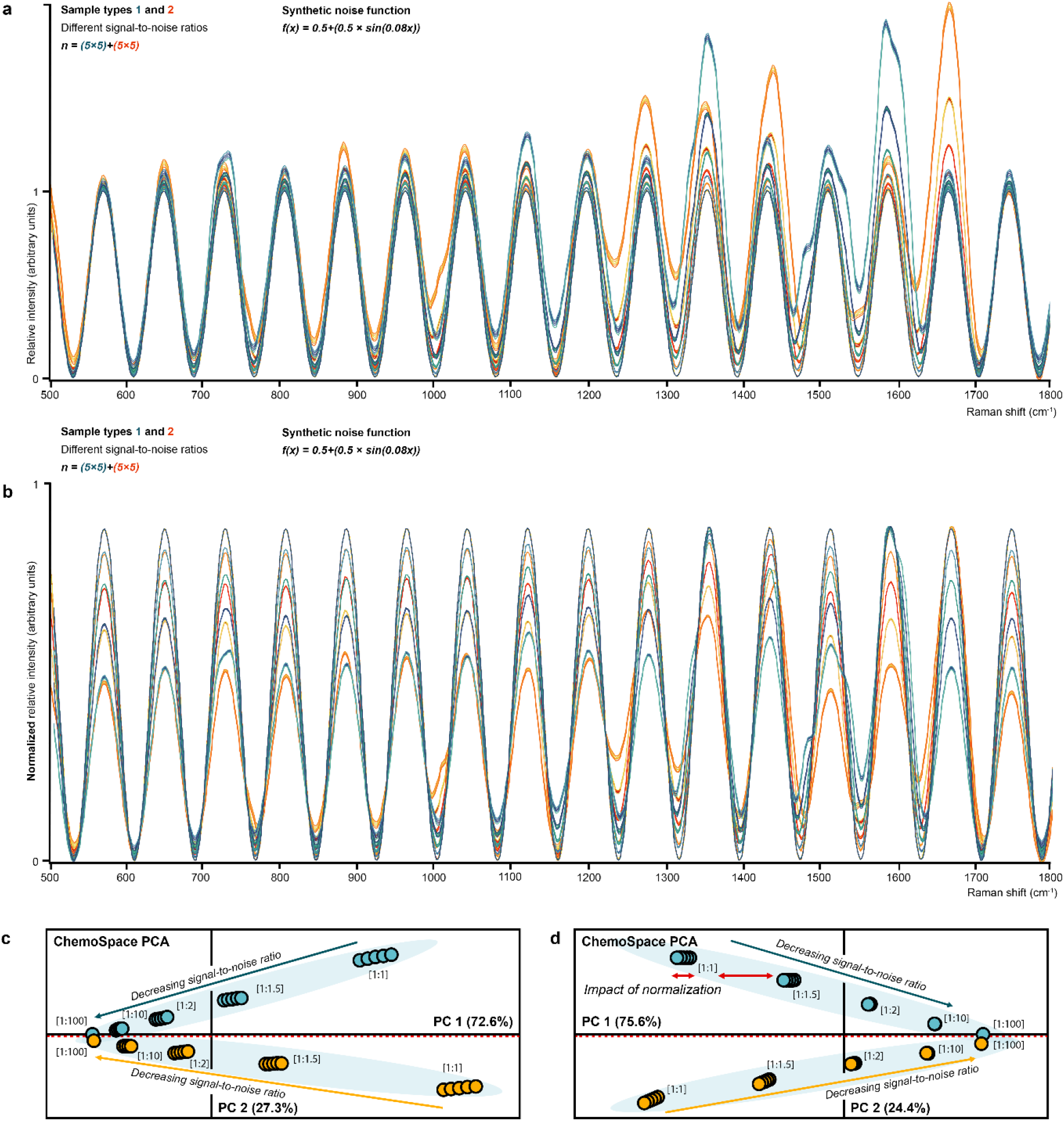
The impact of high-frequency sinusoidal noise and different signal-to-noise ratios on ChemoSpace occupation. **a** Plot of n=5 varieties (synthetic technical replicates) of n=5 differently scaled sets of spectra corresponding to the two sample types (1: teal hues; 2: orange hues), added to the normalized synthetic noise function (for details see figure) over the organic fingerprint region. **b** The same spectra as in **a**, normalized (standard normalization) to the highest peak in each spectrum. **c** ChemoSpace plot across PCs 1 and 2 based on a variance-covariance matrix including select relative intensities (see Methods) extracted from the plot in **a**. Data point fill colors correspond to the sample type (1: teal hues; 2: orange hues; compare **a**). The different signal-to-noise ratios are shown for groups of data points, and the labelled arrows indicate general patterns in the data dristribution across the ChemoSpace. **d** ChemoSpace plot across PCs 1 and 2 based on a variance-covariance matrix including select relative intensities (see Methods) extracted from the plot in **b**. Data point fill colors correspond to the sample type (1: teal hues; 2: orange hues; compare **b**). The different signal-to-noise ratios are highlighted for groups of data points, and the labelled arrows indicate general patterns in the data dristribution across the ChemoSpace.

**Figure 5:**
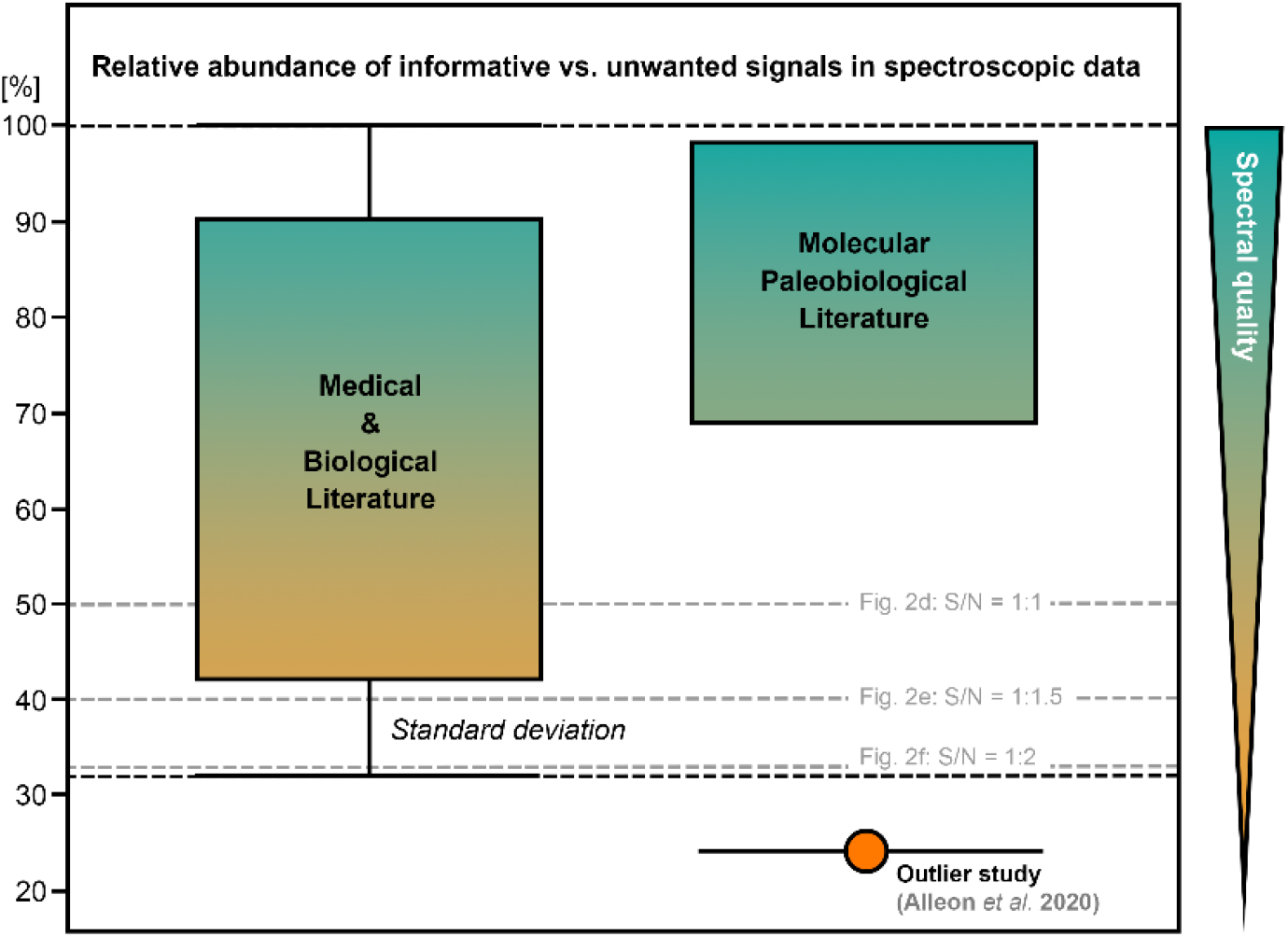
Trends in the relative abundance of informative versus unwanted signals. (compare to the signal-to-noise ratio) in spectroscopic data published in the Molecular Medical and Biological literature (n=8 data sets, n=3-5 replicates were analyzed) and the Molecular Paleobiological literature (n=6 data sets, n=3-5 replicates were analyzed): Categories are separated along the x-axis of the plot). The percentage of true compositional signal relative to the total amount of spectroscopic signal, which includes both compositional and unwanted signals, in the published sets of spectra is shown on the y-axis of the plot. The bars associated with the percentage of informative signal in spectra from Medical and Biological publications represent the standard deviation based on the analyzed spectral sample (±1σ)^27^. For Molecular Paleobiological studies with sufficient spectral data published alongside the article^2,13-14,18,20^, one outlier study was identified^11^. Signal-to-noise ratios (S/N) corresponding to the listed percentage of informative spectral signals in Fig. 2d-f are plotted in form of grey, dashed lines (labelled in the figure). The color gradient in the data bars corresponds to trends in the spectral quality.

**Figure 6:**
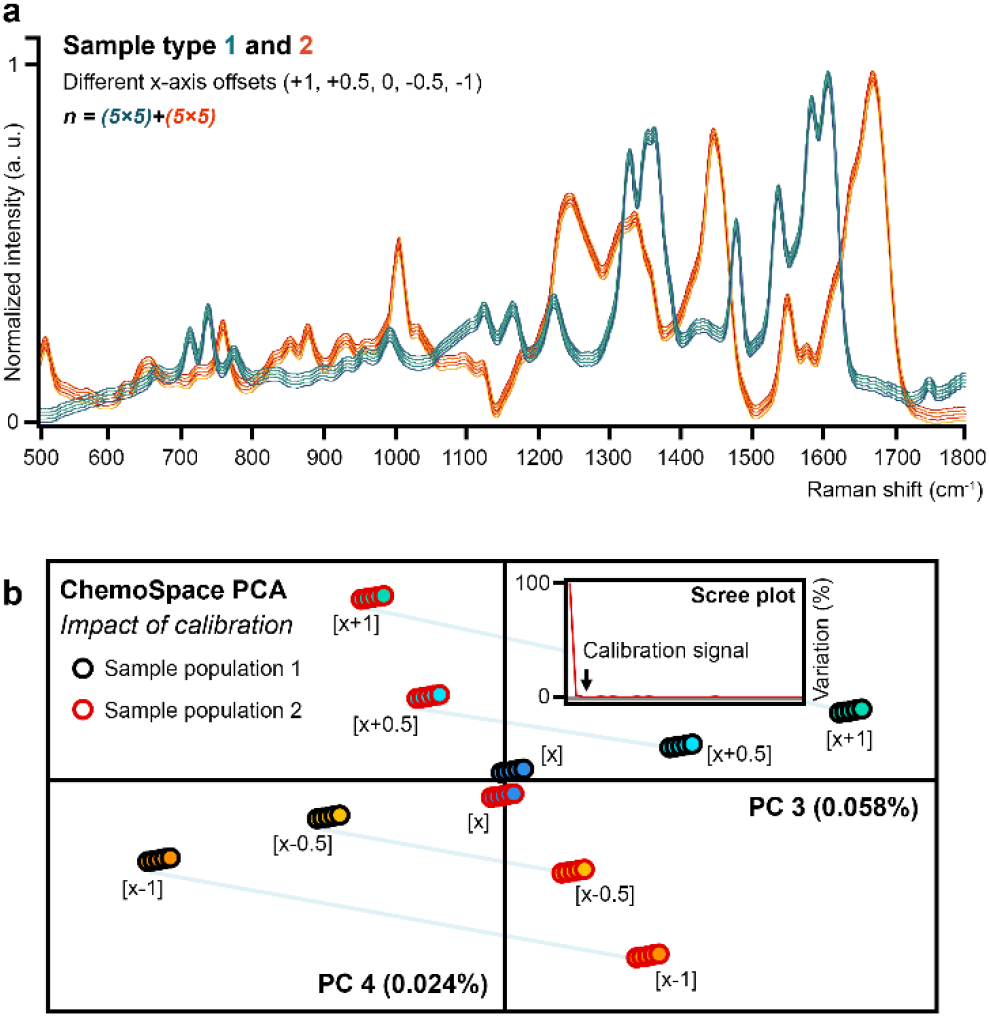
The impact of spectrometer decalibration on the occupation of the ChemoSpace. **a** Plot of n=5 varieties (synthetic technical replicates) of n=5 different x-axis offsets applied to the two sample types (1: teal; 2: orange) over the organic fingerprint region. **b** ChemoSpace across PCs 3 and 4 based on a variance-covariance matrix including select relative intensities (see Methods) extracted from the plot in **a**. Data point outline colors correspond to the sample type (1;2), while fill colors correspond to the x-axis offset (compare **a**). The scree plot of PC loadings indicates the placement of the decalibration signal.

**Figure 7:**
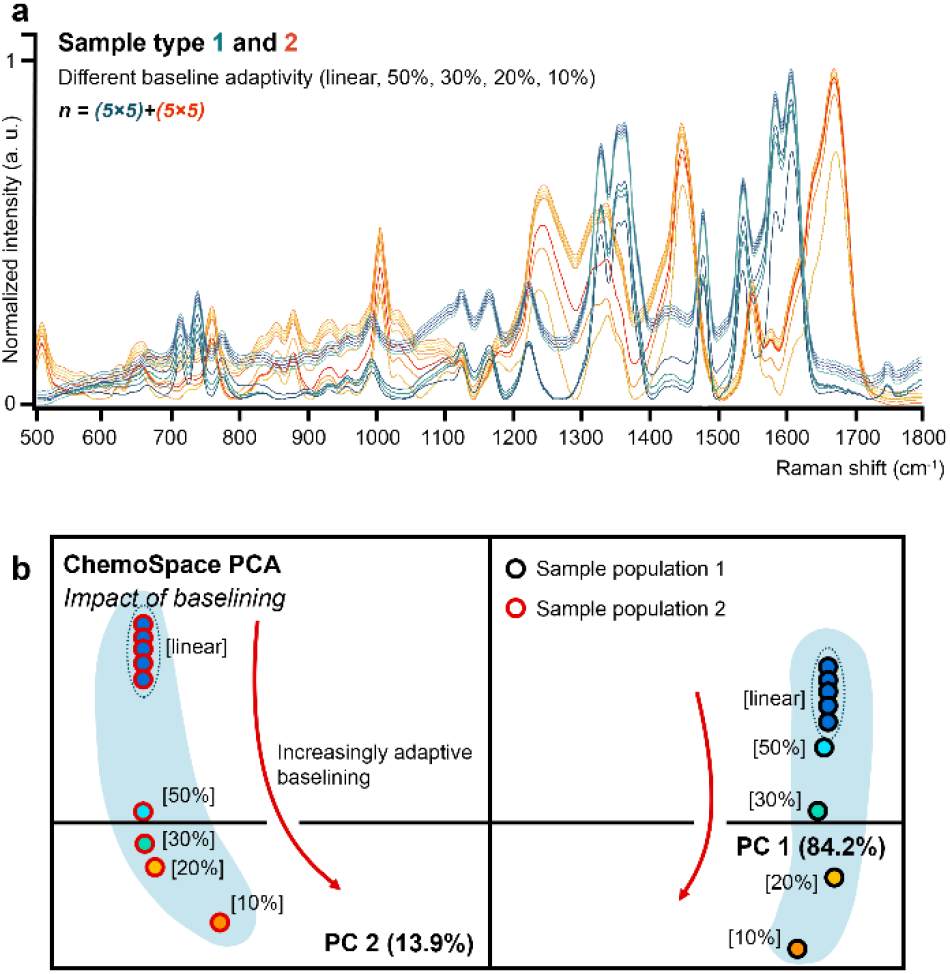
The impact of baseline subtraction on the occupation of the ChemoSpace. **a** Plot of n=5 varieties (synthetic technical replicates) of n=5 different baseline subtraction approaches (in SpectraGryph^26^: linear, 50%, 30%, 20%, 10%) applied to the two sample types (1: teal hues; 2: orange hues) over the organic fingerprint region. **b** ChemoSpace plot across PCs 1 and 2 based on variance-covariance matrix including relative intensities (see Methods) extracted from the spectra in **a**. Data point outline colors correspond to the sample type (1: black; 2: red), while the fill colors correspond to the different baseline subtraction routines (labelled in the figure). Red arrows point towards increasingly adaptive baselines.

## 2. METHODS

In order to illustrate the effects of sample size, instrument features, and spectral processing on ChemoSpace occupation without potential signal distortion, we have selected spectra of two different biological tissues with a simple composition, labelled as sample type 1 and 2. For the purpose of experimentation these spectra were modified in a number of ways: (1) whole-spectral scaling to generate varieties (5–30, depending on the specific analysis) of the same source signal, (2) superimposing synthetic sinusoidal wave functions with broad and narrow frequencies (the latter equal the average Raman band base width in the source spectra), (3) shifting of the x-axis (+1, +0.5, 0, –0.5, –1 cm^-1^ offsets), (4) linear and adaptive baselining (as performed with the SpectraGryph freeware^29^: linear [no offset]; 50%, 30%, 20%, 10% baseline adaptivity options), and (5) spectral normalization relative to the highest peak. All source data are available upon request.

### Impact of the sample: Sample size

Adding samples to a small initial number is expected to result in rotation of the axis separating the two sample types as the amount of variation within the groups increases^22,23^. Stable ChemoSpace occupation requires a representative sample, the number varying depending on the amount of spectral variation in the data set. To determine how many spectra per sample group are required to achieve a stable pattern in ChemoSpace occupation, 30 scaled varieties of the two source spectra (sample type 1 and 2), i.e., a total of 60, were generated. Sets of 3, 5, 6, and 30 spectra per sample type were analyzed and the individual sample sets plotted in SpectraGryph. Relative intensities were extracted from all spectra at the 39 Raman band positions resulting in [3] × [39], [5] × [39], [6] × [39], and [30] × [39] variance-covariance matrices. PC loadings and ChemoSpace plots based on PCs 1 and 2 are shown in Fig. 2.

### Impact of the sample and instrument: Systemic unwanted signals

To determine the impact of systemic unwanted signals^11,12^ on ChemoSpace occupation, the 5 individual varieties of the two source spectra (sample type 1 and 2) were scaled to 100%, ∼66 %, 50%, 10%, 0.1% of their normalized intensity (the highest peak scaled to the value 1) and the results plotted in SpectraGryph (Figs. 3a, b). Two sinusoidal noise functions were computed, one low frequency (*f*(*x*) = 0.5 + (0.5 × sin(0.015*x*))), the other high (*f*(*x*) =0.5 + (0.5 × sin(0.08*x*))) matching the average Raman band base width in the source spectra. The five sets of scaled spectra for sample type 1 and 2 were added to these synthetic noise functions, resulting in five different signal-to-noise ratios: 1:1 (informative signal content: 50%), 1:1.5 (informative signal content: 40%), 1:2 (informative signal content:∼ 33%), 1:10 (informative signal content: ∼ 9%), 1:100 (informative signal content: ∼ 1%). The resulting combined signals for low (all spectral varieties in Fig. 3c, individually scaled sub-samples in Figs. 3d-h) and high (Fig. 4a) frequency noise were plotted separately in SpectraGryph. The sample sets containing low (Fig. 3i) and high (Fig. 4c) frequency noise were subjected to PCA in PAST 3.0, and the resulting PC loadings and ChemoSpaces based on PCs 1 and 2 were extracted.

To contextualize and constrain the signal-to-noise ratios in Raman spectra of modern and fossil biological samples (here subsampled from the literature and plotted under ‘Molecular Paleobiology’), between 3 and 5 technical replicates of organic Raman spectra (as available in the individual studies) published in the fields of medicine and biology were compiled. Technical replicates (Fig. 1a) were plotted in SpectraGryph and whole-spectral data were exported to create corresponding variance-covariance matrices. PC 1 axis loading functions were extracted (Fig. 1c), plotted, and normalized together with one of the 3–5 source spectra in SpectraGryph. Integrals of each spectrum and the corresponding PC 1 axis loading function, which represents the true compositional signal, were calculated over the whole spectral range (resolution differs across published data sets). The area under the PC 1 axis loading function was compared to that under the source spectrum containing the additional unwanted signals, expressed as the percentage of coverage (Fig. 1b). Percentage ranges capturing the relationship between the total spectral signal and the true compositional signal were plotted in PAST 3.0 (Fig. 5). Figure 1 illustrates the process of denoising the biological spectra through PCA.

### Impact of the instrument: Spectrometer decalibration

To characterize how ChemoSpace occupation is impacted by routinely occuring, minute spectrometer decalibration such as might occur during a longer analytical session in response due to changes in room temperature^21^, the 5 scaled varieties of the two source spectra (sample type 1 and 2) were shifted along the x-axis as follows: +1, +0.5, +0, -0.5, -1 cm^-1^, resulting in a total of n=(5 × 5) + (5 × 5)=50 spectral varieties. All resulting spectra were plotted in SpectraGryph (Fig. 6a). A variance-covariance matrix was built based on the intensities of major peaks at 39 band positions: 510, 536, 577, 644, 667, 698, 711, 725, 739, 753, 761, 778, 811, 839, 856, 880, 931, 959, 993, 1005, 1031, 1124, 1165, 1186, 1229, 1249, 1330, 1344, 1356, 1363, 1418, 1445, 1478, 1535, 1550, 1586, 1609, 1676, 1751 cm^-1^ Raman shift. The resulting variance-covariance matrix (50 × 39) was subjected to PCA in PAST 3.0^30^ and the (1) variation explained by the calibration signal, (2) sample separation based on calibration differences along PC 3 and 4 in the Chemospace, and (3) corresponding scree plot are illustrated in Fig. 6b.

### Impact of spectral processing: Spectral baselining

Baseline subtraction is an established approach to the processing of spectra^1^ for increasing comparability when background signals differ across samples as, for example, in sets of spectra from modern and fossil organisms. To capture the influence of baselining on ChemoSpace occupation, 5 varieties of the two source spectra (sample type 1 and 2) were subjected to the linear option and the 50%, 30%, 20%, and 10% adaptive baselining options (no y-axis offset in either case) in SpectraGryph. All n=(5 × 5) + (5 × 5)=50 resulting spectra were plotted in SpectraGryph (Fig. 7a). Excessive (≤ 20% in SpectraGryph) baseline adaptivity leads to partial subtraction of signal associated with the highest peaks in the spectra, and alters the relative signal intensities that encode biosignatures. Relative intensities at the same 39 band positions used to investigate decalibration (above) were extracted from all spectra and incorporated into a [50] x [39] variance-covariance matrix. Principal Component Analysis was performed in PAST 3.0 to capture the impact of different baselines on ChemoSpace occupation reflected in PC loadings and sample position in the ChemoSpace plot based on PCs 1 and 2 (Fig. 7b).

### Impact of spectral processing: Spectral normalization

Normalization scales a spectrum based on the highest peak, a particular selected peak, or the area under the spectral curve^1,29^. It is commonly applied prior to any quantitative analysis^1^ to increase comparability across spectra given the variability of absolute Raman intensities among diverse samples. The combined set of 50 varieties of spectra containing the synthetic, high-frequency noise was plotted (Fig. 3a) to capture the impact of normalization on ChemoSpace occupation. The highest peak of each spectrum was scaled to a value of 1 (the usual approach) using the SpectraGryph normalization option (Fig. 3b). Relative intensities were extracted from all spectra at the 39 Raman shift positions generating a [50] × [39] variance-covariance matrix. Figure 3d shows the resulting PC loadings and ChemoSpace plot based on PCs 1 and 2.

## 3. RESULTS AND DISCUSSION

The effect of sample size, instrument decalibration, and spectral processing on ChemoSpace occupation was simulated in six distinct experiments. Minute changes in spectrometer calibration, the systemic presence of unwanted signals, differences in the spectral signal-to-noise ratios, linear and standard adaptive baseline subtraction, and spectral normalization did not negatively impact the biologically informative grouping of samples in the ChemoSpace. Spectral processing, including baseline correction and normalization prior to PCA, improved data comparability and signal separation. A stable pattern in ChemoSpace occupation is, in this example, reached with as few as 6 spectra per sample type.

### Stable ChemoSpace occupation is reached with less than ten samples

The number of samples required to achieve a stable ChemoSpace occupation is as few as 6 per sample type in this data set (Fig. 2). With 12 samples, the two clusters are separated across PC 2 which accounts for 42.7% of the variance in the data set, while intra-group variance accounts for 57.3% of the total and is captured on PC 1. In contrast to PC loadings, eigenvectors in the ChemoSpace biplot allow the sources of variance in the data, including biological signals within and across tissues, to be differentiated even when cluster separation occurs diagonally in the ChemoSpace. Such eigenvector trajectories allowed us to infer that rotation of the axis separating sample clusters in the ChemoSpace results from an increase in the contribution of intra-group variability to the total variance as spectra are added: Intra-group variation becomes the primary source of variance and is displayed along PC 1. The sampling strategy should reflect the question addressed. In integrative data sets including modern and fossil tissues, ChemoSpace grouping will account more accurately for variation in different modes of (diagenetic) alteration of a biological tissue, for example, when an increasing number of fossil samples from different depositional settings is considered.

### Principal Component Analysis eliminates systemic unwanted signals

PCA is based on a variance-covariance matrix. The focus on covariance rather than absolute spectral differences (Fig. 3c) allows unwanted signals, such as sample- or instrument-related spectral features^11,12^, to be eliminated. In addition, the extraction of relative intensities at informative wavenumber positions allows the variation relevant to a given question to be emphasized^24^. An omnipresent signal cannot be a primary source of variance in the data. PCA separates the two clusters corresponding to signals 1 and 2, regardless of the frequency of a periodic, sytemically present, unwanted signal, or the total spectral signal-to-noise ratio (Figs. 3i, 4c). Although decreasing the signal-to-noise ratio results in convergence of the two sample clusters in ChemoSpace, they are clearly separated even with a signal-to-noise ratio of 1:100 (Figs. 3h-i, 4c). The spectra modelled here (with signal-to-noise ratios ranging from 1:100 to 1:1) include a higher amount of unwanted signal than that in most published spectra. Informative spectral content ranges from ∼42 – 90% (± 1σ) in the biological and medical literature (Fig. 5, based on 8 spectral data sets, 3-5 replicates were analyzed), and ∼69 – 98% in the molecular paleobiological literature^2,12-16,18-20^ excluding one statistical outlier^11^ (Fig. 5, based on 6 spectral data sets, 3-5 replicates were analyzed). Most of the ranges of compositional signal content in the published spectra overlap. Comparatively high signal-to-noise content in carbonaceous fossilization products is the result of smoother textures following dehydration and reduced fluorescence (Fig. 5). Regardless of the type of unwanted signal present in biological Raman spectra, PCA reliably extracts informative features (see Fig. 1 for denoising).

### Minor in-session spectrometer decalibration does not impact ChemoSpace biosignatures

Spectrometer decalibration accounting for a ±1 cm^−1^ shift in wavenumber is only evident across PC 3 and 4 (Fig. 6), and explains less than 0.1% variance in this data set. Thus any type of biosignature accounting for more than 0.06% variance (loading PC 3) in the data set will outweigh the decalibration signal in the ChemoSpace. Previously published spectroscopic biosignatures^5-7,13-20^ exceed the amount of variance resulting from decalibration by at least two orders of magnitude.

### Standard adaptive baselining increases comparability and ChemoSpace signal extraction

It is essential to subtract spectral backgrounds without affecting informative bands to prevent background differences appearing as major variance and overprinting biosignatures^1^. Linear baseline subtraction (Fig. 7a) does not completely remove non-linear background signals, which are common in often heterogenous and stratified spectra of biological tissues, and may introduce or amplify spectral incomparability (see linear baseline substraction applied to sample spectrum 1 in Fig. 7a). Adaptive baselining, in contrast, eliminates all background signal (Fig. 7a) regardless of shape. Baselining may result in minor spatial convergence of informative clusters (sample type 1 and 2) in the ChemoSpace (Fig. 7b) if adaptivity exceeds the standard (less of the original spectral signal remains; treshold determined here: <30% in SpectraGryph^26^): As baseline adaptivity increases beyond the standard, broad Raman bands of high intensity lose comparatively more signal than narrow bands with relatively low intensity (Fig. 7a). This loss of the biological signal encoded in informative band ratios decreases the separation of groups in the ChemoSpace (indicated by the red arrows in Fig. 7b). Standard adaptive baselines (treshold: ≥ 30% in SpectraGryph) increase intra-group comparability without cluster convergence, resulting in the collapse of individual spectral data points within a sample type in the ChemoSpace (Fig. 7b).

### Normalization increases comparability

Spectral normalization emphasizes key differences within a sample set by amplifying scaled variation in spectrum intensity (Fig. 4b). The ChemoSpace PCA shows how normalization increases direct comparability (all spectra share a highest peak scaled to 1, rather than ranging in intensity counts over orders of magnitude) across synthetic replicates as demonstrated by the closer grouping of data points within a subsample (Fig. 4d). Normalization also homogenizes the distribution of spectral subsamples (with different S/N ratios) within clusters associated with sample signals 1 and 2 – an inference based on resulting uniform spacing of data point groups subsampled by S/N ratio (Fig. 4d), compared to non-uniform data group spacing among non-normalized spectra (Fig. 4c). Most biosignatures are encoded in the relative abundance of functional groups^1,2^ so spectral normalization (based on the highest peak in the spectrum, a different informative peak, or a spectral area) facilitates the extraction of a meaningfully comparable signal. The mode of normalization applied depends on the specific question, and the nature and comparability of intensities in the sample set.

## CONCLUSIONS

Quantification of the impact of sample size, instrument features, and spectral processing on the occupation of ChemoSpace provides an analytical framework for the extraction of molecular biosignatures from spectroscopic fingerprints of tissues from extant and extinct organisms: Minor instrument decalibration during an analytical session does not overprint key biological signatures in a ChemoSpace PCA. Spectral processing routines, such as standard adaptive baseline subtraction, as well as normalization prior to statistical analysis of spectra, increase data comparability and extraction. Stable ChemoSpace occupation can be achieved with fewer than ten spectra per sample group when analyzing biosignatures. PCA eliminates systemic unwanted signals, regardless of waveform, periodicity, frequency or amplitude, and groups samples in the ChemoSpace by informative property, even at relatively low signal-to-noise ratios. The ChemoSpace approach to biosignatures represents a powerful tool for exploring, denoising, and integrating molecular biological information from modern and ancient organismal samples.

## ACKNOWLEDGEMENTS

The authors thank D. Briggs for helpful comments and edits, and J. Eiler, M. Brown, and G. Rossman for helpful conversations. We acknoweldge funding from the Agouron Geobiology Institute (JW) and the TAWANI Foundation (PRH).

## REFERENCES

1. Butler, H.J., Ashton, L., Bird, B., Cinque, G., Curtis, K., Dorney, J., Esmonde-White, K., Fullwood, N.J., Gardner, B., Martin-Hirsch, P.L. and Walsh, M.J., 2016. Using Raman spectroscopy to characterize biological materials. Nature protocols, 11(4), pp. 664–687.

2. Wiemann, J., Crawford, J.M. and Briggs, D.E., 2020. Phylogenetic and physiological signals in metazoan fossil biomolecules. Science Advances, 6(28), p. eaba6883.

3. Talari, A.C.S., Movasaghi, Z., Rehman, S. and Rehman, I.U., 2015. Raman spectroscopy of biological tissues. Applied spectroscopy reviews, 50(1), pp. 46–111.

4. Manoharan, R., Wang, Y. and Feld, M.S., 1996. Histochemical analysis of biological tissues using Raman spectroscopy. Spectrochimica Acta Part A: Molecular and Biomolecular Spectroscopy, 52(2), pp. 215–249.

5. Mahadevan-Jansen, A. and Richards-Kortum, R.R., 1996. Raman spectroscopy for the detection of cancers and precancers. Journal of biomedical optics, 1(1), pp. 31–70.

6. Talari, A.C.S., Evans, C.A., Holen, I., Coleman, R.E. and Rehman, I.U., 2015. Raman spectroscopic analysis differentiates between breast cancer cell lines. Journal of Raman Spectroscopy, 46(5), pp. 421–427.

7. Li, X., Yang, T., Li, S., Wang, D., Song, Y. and Zhang, S., 2016. Raman spectroscopy combined with principal component analysis and k nearest neighbour analysis for non-invasive detection of colon cancer. Laser Physics, 26(3), p. 035702.

8. Pasteris, J.D. and Beyssac, O., 2020. Welcome to Raman spectroscopy: successes, challenges, and pitfalls. Elements, 16(2), pp. 87–92.

9. Olcott Marshall, A. and Marshall, C.P., 2015. Vibrational spectroscopy of fossils. Palaeontology, 58(2), pp. 201–211.

10. Schopf, J.W., Kudryavtsev, A.B., Agresti, D.G., Wdowiak, T.J. and Czaja, A.D., 2002. Laser–Raman imagery of Earth’s earliest fossils. Nature, 416(6876), pp. 73–76.

11. Alleon, J., Montagnac, G., Reynard, B., Brulé, T., Thoury, M. and Gueriau, P., 2021. Pushing Raman spectroscopy over the edge: purported signatures of organic molecules in fossil animals are instrumental artefacts. BioEssays, 43(4), p. 2000295.

12. Wiemann, J. and Briggs, D.E., 2022. Raman spectroscopy is a powerful tool in molecular paleobiology: An analytical response to Alleon et al. (https://doi. org/10.1002/bies.202000295). BioEssays, 44(2), p. 2100070.

13. Norell, M.A., Wiemann, J., Fabbri, M., Yu, C., Marsicano, C.A., Moore-Nall, A., Varricchio, D.J., Pol, D. and Zelenitsky, D.K., 2020. The first dinosaur egg was soft. Nature, 583(7816), pp. 406–410.

14. McCoy, V.E., Wiemann, J., Lamsdell, J.C., Whalen, C.D., Lidgard, S., Mayer, P., Petermann, H. and Briggs, D.E., 2020. Chemical signatures of soft tissues distinguish between vertebrates and invertebrates from the Carboniferous Mazon Creek Lagerstätte of Illinois. Geobiology, 18(5), pp. 560–565.

15. Tripp, M., Wiemann, J., Brocks, J., Mayer, P., Schwark, L. and Grice, K., 2022. Fossil Biomarkers and Biosignatures Preserved in Coprolites Reveal Carnivorous Diets in the Carboniferous Mazon Creek Ecosystem. Biology, 11(9), p. 1289.

16. Fabbri, M., Wiemann, J., Manucci, F. and Briggs, D.E., 2020. Three-dimensional soft tissue preservation revealed in the skin of a non-avian dinosaur. Palaeontology, 63(2), pp. 185–193.

17. Loron, C.C., Rodriguez Dzul, E., Orr, P.J., Gromov, A.V., Fraser, N.C. and McMahon, S., 2023. Molecular fingerprints resolve affinities of Rhynie chert organic fossils. Nature Communications, 14(1), p. 1387.

18. Wiemann, J., Yang, T.R. and Norell, M.A., 2018. Dinosaur egg colour had a single evolutionary origin. Nature, 563(7732), pp. 555–558.

19. Wiemann, J., Menéndez, I., Crawford, J.M., Fabbri, M., Gauthier, J.A., Hull, P.M., Norell, M.A. and Briggs, D.E., 2022. Fossil biomolecules reveal an avian metabolism in the ancestral dinosaur. Nature, 606(7914), pp. 522–526.

20. Wiemann, J., Fabbri, M., Yang, T.R., Stein, K., Sander, P.M., Norell, M.A. and Briggs, D.E., 2018. Fossilization transforms vertebrate hard tissue proteins into N-heterocyclic polymers. Nature Communications, 9(1), p. 4741.

21. Hutsebaut, D., Vandenabeele, P. and Moens, L., 2005. Evaluation of an accurate calibration and spectral standardization procedure for Raman spectroscopy. Analyst, 130(8), pp. 1204–1214.

22. Abdi, H. and Williams, L.J., 2010. Principal component analysis. Wiley interdisciplinary reviews: computational statistics, 2(4), pp. 433–459.

23. Zelditch, M.L., Swiderski, D.L. and Sheets, H.D., 2012. Geometric morphometrics for biologists: a primer. Academic press.

24. Lewis, P.D. and Menzies, G.E., 2015. Vibrational spectra, principal components analysis and the horseshoe effect. Vibrational Spectroscopy, 81, pp. 62–67.

25. Lee, W., Lenferink, A.T., Otto, C. and Offerhaus, H.L., 2020. Classifying Raman spectra of extracellular vesicles based on convolutional neural networks for prostate cancer detection. Journal of raman spectroscopy, 51(2), pp. 293–300.

26. Sattlecker, M., Stone, N., Smith, J. and Bessant, C., 2011. Assessment of robustness and transferability of classification models built for cancer diagnostics using Raman spectroscopy. Journal of Raman spectroscopy, 42(5), pp. 897–903.

27. Morais, C.L., Lima, K.M., Singh, M. and Martin, F.L., 2020. Tutorial: multivariate classification for vibrational spectroscopy in biological samples. Nature Protocols, 15(7), pp. 2143–2162.

28. Beyssac, O., 2020. New trends in Raman spectroscopy: from high-resolution geochemistry to planetary exploration. Elements: An International Magazine of Mineralogy, Geochemistry, and Petrology, 16(2), pp. 117–122.

29. Menges, F., 2020. Spectragryph Optical Spectroscopy Software, Version 1.2. 14. Oberstdorf, Germany 2019. Available online: https://www.effemm2.de/spectragryph/

30. Hammer, Ø., Harper, D.A. and Ryan, P.D., 2001. PAST: Paleontological statistics software package for education and data analysis. Palaeontologia electronica, 4(1), p. 9.

